# A Collagen-Based 3D Microtissue Platform Preserves Structural, Molecular, and Therapeutic Heterogeneity in Ovarian Cancer

**DOI:** 10.64898/2026.10.25.761684

**Authors:** Parnaz Soori, Mohammadreza Javan, Lacey Winstone, Breanne Bevelander, Raquel Santamaria Cuevas, M. Dean Chamberlain

## Abstract

**Background:** Ovarian cancer is comprised of biologically diverse tumour histotypes that are difficult to represent using conventional two-dimensional (2D) culture. These systems lack the tissue organization and extracellular matrix interactions found in native tumours. We developed a type I collagen-based three-dimensional (3D) microtissue model to determine whether a tumor supportive matrix environment could maintain key characteristics of different ovarian cancer histotypes.

**Methods:** 3D collagen microtissues were generated using five ovarian cancer cell lines: OVCAR3, SNU119, KURAMOCHI, OVTOKO, and SKOV3. Long-term growth, viability, and morphology were assessed using quantitative growth analysis, confocal microscopy, and longitudinal bright-field imaging. Tumour architecture and histotype-associated phenotypes were evaluated by hematoxylin and eosin (H&E) staining and immunohistochemistry (IHC). Therapeutic responses to carboplatin, paclitaxel, olaparib, and niraparib were compared between 3D microtissues and corresponding 2D monolayer cultures.

**Results:** All five models remained viable and showed sustained growth within the collagen scaffold, supported by growth-curve analysis and confocal imaging. Despite being maintained in the same matrix environment, the models developed distinct 3D architectures. OVCAR3, KURAMOCHI and SNU119 formed prominent papillary-like structures of high-grade serous carcinoma (HGSC), while OVTOKO and SKOV3 displayed distinct surface-associated growth patterns of clear cell and endometrioid ovarian carcinomas, respectively. H&E analysis confirmed these architectural differences, and IHC profiling demonstrated preservation of selected histotype-associated features, including distinct PAX8, WT1, and p53 expression patterns. Therapeutic responses varied across cell lines and treatments, with 3D microtissues generally maintaining greater viability following treatment than corresponding 2D cultures. These findings indicate that the collagen environment supports long-term tumour growth while preserving ovarian cancer histotype specific structural, molecular, and therapeutic heterogeneity.

**Conclusions:** The collagen-based microtissue platform preserved structural, molecular, and therapeutic heterogeneity across distinct ovarian cancer models within a shared 3D environment. By maintaining tumour-associated architecture, distinct immunophenotypic features, and model-specific therapeutic responses, the platform captures key aspects of ovarian cancer biology that are not adequately represented in conventional 2D culture. These findings support its use as a physiologically relevant system for investigating ovarian cancer heterogeneity, tumour–matrix interactions, and therapeutic response.

## 1. Introduction

Ovarian cancer is a highly heterogeneous disease that includes several histological and molecular histotypes. These histotypes differ in their biological behaviour, molecular characteristics, and response to treatment ^(1–3)^. High-grade serous carcinoma (HGSC) is the most common and lethal epithelial ovarian cancer, while clear-cell, endometrioid, low-grade serous, and mucinous carcinomas represent distinct disease entities ^(4)^. Substantial heterogeneity also exists within individual ovarian-cancer histotypes, contributing to variation in tumour progression, treatment response, and the emergence of drug resistance ^(5)^. Accordingly, in vitro models that capture clinically relevant aspects of this heterogeneity are valuable for investigating tumour biology and evaluating therapeutic strategies ^(6, 7)^.

Conventional two-dimensional (2D) cultures remain widely used in ovarian cancer research. However, these systems reproduce only a limited aspect of the tumour environment. Cells grown on rigid plastic surfaces adopt an artificial planar organization and lack many of the extracellular matrix (ECM) interactions present in intact tumours ^(8, 9)^. As a result, 2D culture can alter cell morphology, proliferation, signalling, gene expression, and response to therapy ^(9, 10)^. Drug responses measured under these conditions may therefore differ from those occurring within a three-dimensional (3D) tumour environment ^(11)^. This is particularly important in ovarian cancer, where interactions between tumour cells and the surrounding ECM influence disease progression and therapeutic resistance ^(12)^. Type I collagen is a major component of the tumour ECM and plays an active role in cancer biology. In addition to providing structural support, collagen regulates cell adhesion, migration, proliferation, survival, and mechanotransduction ^(13–15)^. Changes in collagen abundance, organization, and mechanical properties are also associated with tumour progression and treatment resistance ^(15–17)^. Collagen-based 3D systems can therefore provide a more physiologically relevant environment for studying ovarian cancer cells. Importantly, they restore tumour–matrix interactions that are largely absent from conventional monolayer cultures ^(18)^.

Several 3D approaches have been developed to improve ovarian cancer modelling, including spheroids, organoids, hydrogels, and scaffold-based systems. These models can reproduce aspects of tumour organization and often generate therapeutic responses that differ from 2D culture ^(10, 11)^. However, each approach has limitations. Scaffold-free spheroids primarily reproduce cell–cell interactions and provide limited control over cell–matrix interactions ^(19)^. Matrigel-based 3D models provide a biologically active-matrix environment but are limited by their complex and incompletely defined composition and potential batch-to-batch variability, which may affect experimental reproducibility ^(20, 21)^. Beyond these limitations, modelling surface-associated tumour growth is relevant to ovarian cancer, which commonly disseminates within the peritoneal cavity ^(22)^. Disseminated cells and multicellular aggregates can adhere to mesothelial surfaces and form deposits on the peritoneum, omentum, and other abdominal organs ^(23)^. Scaffold-free spheroids and fully embedded matrix systems model other important aspects of ovarian cancer biology but may not directly model tumour growth along an accessible matrix surface ^(22–24)^. Collagen-based scaffolds with an exposed surface may therefore provide a versatile platform for studying tumour-cell attachment, surface-associated expansion, and multicellular organization ^(25)^.

A comprehensive evaluation of 3D tumour models should extend beyond tumour growth or morphology alone. It should consider features across multiple levels of tumour biology ^(19)^. These include long-term growth, tissue architecture, histotype-associated molecular characteristics, and heterogeneous responses to therapy ^(26, 27)^. Evaluating these properties within the same matrix environment is particularly valuable. It allows differences between tumour models to emerge under controlled conditions and provides a means of examining how intrinsic tumour characteristics interact with the extracellular matrix ^(28)^.

In this study, we developed a type I collagen-based 3D microtissue platform using five ovarian cancer cell lines with different histological backgrounds. OVCAR3, SNU119, and KURAMOCHI were used as serous/HGSC-associated models, OVTOKO as an ovarian clear-cell carcinoma model, and SKOV3 as endometrioid ovarian carcinoma model. We first evaluated long-term viability and surface-associated tumour organization on the collagen scaffold. Morphological and histological analyses were then used to characterize differences in tumour architecture. Immunohistochemistry (IHC) was used to determine whether histotype-associated phenotypes were maintained during prolonged 3D culture. Finally, functional responses were evaluated using carboplatin, paclitaxel, olaparib, and niraparib and compared with corresponding 2D cultures. Together, these analyses were designed to determine whether a common collagen microenvironment could preserve the structural, molecular, and functional heterogeneity of ovarian cancer within a single experimentally tractable platform.

## 2. Materials and Methods

### 2.1 Drugs and Reagents

Stock solutions of Carboplatin (ApexBio Technology, Cat. No. A2171) were prepared in sterile water. Paclitaxel (Cayman Chemical, Cat. No. 10461), Niraparib (AdooQ Bioscience, Cat. No. A11026), and Olaparib (Ann Arbor, Cat. No. 10621) were dissolved in dimethyl sulfoxide (DMSO; BioBasic, Cat. No. BP231-100). Following preparation, all drug stocks were aliquoted and stored at −80 °C until further use.

### 2.2 Two-Dimensional Cell Culture

Five ovarian cancer cell lines were used in this study and cultured according to American Type Culture Collection (ATCC) guidelines, with minor modifications where specified. OVTOKO cells were maintained in RPMI-1640 medium (Gibco, Cat. No. 11875-093) supplemented with 10% fetal bovine serum (FBS; Corning, Cat. No. 35-010-CV) and 1% penicillin-streptomycin (Gibco, Cat. No. 15140-122). OVCAR3 cells were cultured in Advanced DMEM/F-12 medium (Gibco, Cat. No. 12634-010) containing 20% FBS, 1% penicillin-streptomycin, 1% L-glutamine (Corning, Cat. No. 25-005-CI), and 0.01 mg/mL insulin (Gibco, Cat. No. 12585-014). SNU119, SKOV3, and KURAMOCHI cells were maintained in Advanced DMEM/F-12 medium supplemented with 10% FBS, 1% penicillin-streptomycin, and 1% L-glutamine. All cell lines were cultured at 37°C in a humidified atmosphere containing 5% CO₂.

### 2.3 Microtissue Fabrication and Culture

Collagen microtissues with OVCAR3, SNU119, KURAMOCHI, OVTOKO, and SKOV3 ovarian cancer cells grown on their surface were generated using a modified collagen-casting technique ^(29)^. Briefly, type I bovine collagen (3 mg/mL; PureCol®, Cat. No. 5005) was mixed with 10× minimum essential medium (MEM; Sigma-Aldrich, Cat. No. M9288) to achieve a final concentration of 1× MEM. The collagen solution was neutralized to pH 7.4 using 1 M sodium bicarbonate and maintained on ice to prevent premature gelation.

The neutralized collagen mixture was loaded into sterile polyethylene tubing (PE90; BD Biosciences, Cat. No. 14117012D) and incubated at 37°C in a humidified atmosphere containing 5% CO₂ for 1 hour to allow complete polymerization. Following gelation, the collagen-filled tubing was cut into uniform 2-mm segments using a custom-designed 3D-printed cutting device which guided a razor blade through parallel slots spaced at fixed intervals. The resulting collagen microtissues were collected in 50-mL conical tubes containing culture medium and gently vortexed to release the microtissues from the plastic and collected in a new 50 mL conical tube. For each 1.2-m length of collagen-filled tubing, cells harvested from two T-75 flasks at approximately 80% confluence were seeded onto the collagen surface. Harvested ovarian cancer cells were then seeded onto the surface of the preformed collagen microtissues to promote surface-associated cellular attachment and organization. Following fabrication and cell seeding, microtissues were maintained in non-tissue-culture-treated 10-cm Petri dishes (Thermo, Cat. No. 150460) under cell line-specific culture conditions. SNU119, KURAMOCHI, OVTOKO, and SKOV3 microtissues were maintained in a 1:1 (v/v) mixture of Advanced DMEM/F-12 medium supplemented with 10% fetal bovine serum (FBS), 1% penicillin-streptomycin, and 1% L-glutamine, and endothelial growth medium (EGM-2; Lonza, Cat. No. CC-3162). OVCAR3 microtissues were cultured in a 1:1 (v/v) mixture of EGM-2 and Advanced DMEM/F-12 supplemented with 20% FBS, 1% penicillin-streptomycin, 1% L-glutamine, and 0.01 mg/mL insulin.

### 2.4 Microtissue Viability Assessment

Microtissue viability was assessed using the CellTiter-Glo® 3D Cell Viability Assay (Promega, Cat. No. G9681), which measures ATP as an indicator of metabolically active viable cells. Microtissues from each ovarian cancer cell line were evaluated on days 7, 14, and 21 of culture. At each time point, three microtissues from each cell line were transferred to individual wells of a 96-well plate and processed according to the manufacturer’s protocol. CellTiter-Glo® 3D reagent was added to each well, and the contents were mixed to promote microtissue disruption and ATP release. Following incubation, luminescence was measured using a microplate reader. Background luminescence from medium-only controls was subtracted from each measurement. Luminescence values were compared across time points as a relative measure of metabolic viability throughout the 21-day culture period.

### 2.5 Confocal viability imaging

Microtissue viability was further assessed by confocal microscopy on days 14, and 21 of culture. Microtissues were stained with calcein-AM to identify viable cells and ethidium homodimer-1 (EthD-1) to identify membrane-compromised, non-viable cells. Following staining, microtissues were imaged using a Zeiss LSM 900 confocal laser scanning microscope (Carl Zeiss Microscopy). Optical sections were acquired through the microtissues using excitation and emission settings appropriate for each fluorophore. Confocal imaging was used to visualize the spatial distribution of viable and non-viable cells within the 3D microtissues and to complement the longitudinal metabolic viability measurements.

### 2.6 Histology

Microtissues were rinsed in phosphate-buffered saline (PBS) and fixed overnight in 10% neutral buffered formalin (NBF). Following fixation, samples were embedded in 1.5% agarose using cryomolds to facilitate downstream processing. Agarose blocks were subsequently post-fixed in 10% NBF for 24 h, followed by two washes in 70% ethanol, and stored in 70% ethanol until processing. After routine tissue processing and sectioning, slides were stained with hematoxylin and eosin (H&E). Whole-slide imaging was performed using an Aperio Scanscope digital slide scanner.

### 2.7 Immunohistochemistry (IHC)

### 2.8 Drug screening assay

Drug response studies were conducted in both 3D collagen microtissues and conventional 2D monolayer cultures. For 3D assays, microtissues were cultured for 7 days before treatment. Subsequently, three microtissues suspended in 100 μL of complete culture medium were transferred to individual wells of non-tissue-culture-treated 96-well plates. For 2D experiments, cells were seeded in tissue-culture-treated 96-well plates (Corning, Cat. No. CLS353376) at a density of 2,000 cells per well in 100 μL of complete medium and allowed to attach for 24 h before treatment.

Drug stocks were diluted in culture medium to generate a seven-point serial dilution of the drug series at twice the desired final concentrations. Equal volumes (100 μL) of each drug dilution were added to the wells to achieve the target treatment concentrations. All experimental conditions were performed in triplicate. Controls included medium-only wells and vehicle-control wells containing DMSO or water at concentrations matching those used in the highest drug treatment groups.

Following drug addition, plates were incubated for 4 days at 37°C in a humidified atmosphere containing 5% CO₂. Cell viability was quantified after the 4 days of treatment using the CellTiter-Glo® 3D Cell Viability Assay according to the manufacturer’s protocol. Luminescence readings were normalized to the average signal obtained from medium-only control wells.

### 2.9 Brightfield Microscopy

Brightfield imaging was performed to assess microtissue morphology at defined time points. Microtissues were collected on days 14 and 21, and a few intact microtissues were transferred directly onto glass microscope slides. Imaging was performed immediately after placement to prevent dehydration. Brightfield images were acquired using an upright microscope equipped with a Moticam X5 Plus digital camera (Motic) mounted via a C-mount adaptor (Motic, part no. 9126021). Images were captured using the same acquisition settings across all samples.

### 2.10 Statistical analysis

Quantitative data were analyzed and visualized using GraphPad Prism 11 (GraphPad Software, San Diego, CA, USA). Drug response curves for carboplatin, paclitaxel, olaparib, and niraparib were generated by fitting normalized cell viability data using nonlinear regression with a variable-slope four-parameter logistic (4PL) model [log(inhibitor) versus response]. The model estimates the upper and lower response plateaus, the Hill slope, and the half-maximal inhibitory concentration (IC₅₀). IC₅₀ values derived from the fitted curves were used to compare drug sensitivity among ovarian cancer cell lines and between 2D and 3D culture models.

## 3. Results

### 3.1 Collagen microtissues support long-term growth and viability of diverse ovarian cancer cell lines

To evaluate the capacity of the collagen-based platform to support sustained 3D culture, microtissues were established using OVCAR3, SNU119, KURAMOCHI, OVTOKO, and SKOV3 cells and maintained for 21 days. Brightfield imaging revealed distinct cell line-dependent growth patterns at days 14 and 21 (Fig. 1A). OVCAR3 and SNU119 developed irregular microtissue contours with prominent surface projections, whereas KURAMOCHI exhibited a more heterogeneous pattern of cellular organization along the collagen scaffold. Unlike OVCAR3 and SNU119, which appeared to form more continuous cellular coverage of the collagen surface, KURAMOCHI displayed regions of incomplete surface coverage that remained visible on day 21. Nevertheless, all three serous-associated models exhibited finger-like projections extending from the microtissue surface, resembling aspects of the papillary architecture associated with high-grade serous ovarian carcinoma. In contrast, OVTOKO exhibited a relatively compact morphology with a smoother and more uniform outer contour and fewer prominent surface projections. SKOV3 also displayed a comparatively compact overall morphology but showed more extensive cellular accumulation and a less uniform peripheral contour than OVTOKO (Fig. 1A).

**Figure 1.**
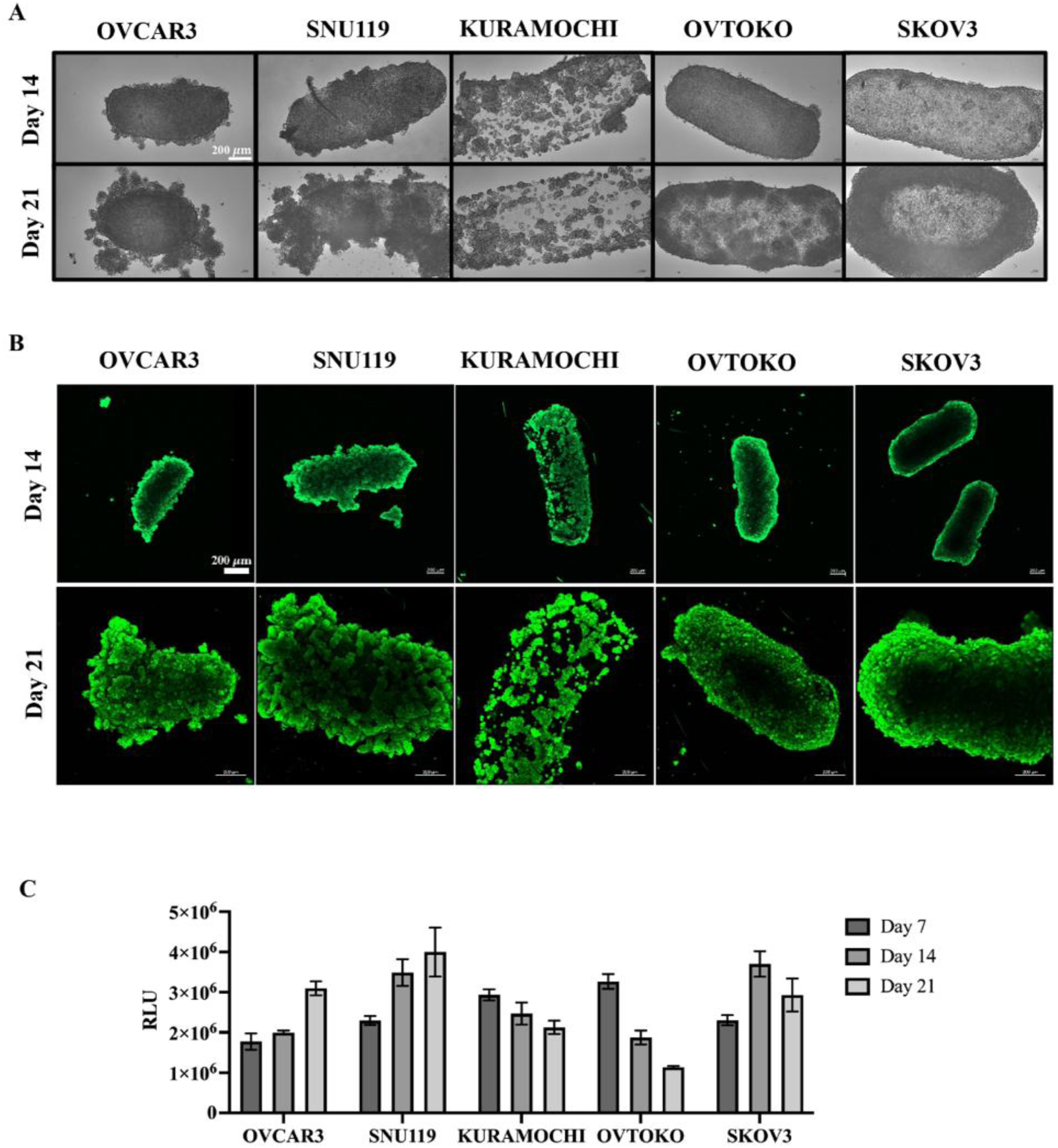
Morphological characterization and long-term viability of ovarian cancer microtissues in a 3D collagen-based culture system. **(A)** Representative bright-field images showing the morphological organization of five ovarian cancer cell lines (OVCAR3, SNU119, KURAMOCHI, OVTOKO, and SKOV3) cultured on collagen-based microtissues at days 14 and 21. Bright-field images were acquired at 4× magnification. Images illustrate cell line-dependent differences in microtissue morphology and cellular organization. Scale bar = 200mm. **(B)** Representative merged confocal fluorescence images showing viable cells stained with calcein-AM (green) and EthD-1 at days 14 and 21. Confocal images were acquired at 2.5X magnification on day 14 (top row) and 5X magnification on day 21 (bottom row). Scale bar = 200mm. **(C)** Longitudinal assessment of cellular metabolic viability using the CellTiter-Glo 3D assay at days 7, 14, and 21. Results are expressed as relative luminescence units (RLU), reflecting ATP-associated metabolic activity across the five ovarian cancer models.

Confocal imaging live/dead staining demonstrated viable-cell coverage across all five models at days 14 and 21 (Fig. 1B). By day 21, an apparent increase in the extent of calcein-AM-positive cellular coverage was observed across the microtissues, although the spatial distribution of viable cells varied among the models. Longitudinal ATP-based viability measurements revealed distinct metabolic activity profiles across the five cell lines (Fig. 1C). OVCAR3 and SNU119 exhibited increasing raw luminescent signals over time, whereas KURAMOCHI and OVTOKO showed declining signals following day 7. SKOV3 exhibited an increase at day 14 followed by a reduction at day 21. Notably, the declining ATP signals in selected models contrasted with the apparent increase in viable-cell coverage observed by confocal imaging. Nevertheless, detectable metabolic activity was maintained across all five models at day 21. These findings demonstrate that the collagen-based platform supports long-term maintenance of ovarian cancer microtissues while revealing cell line-dependent differences in morphology, viable-cell distribution, and metabolic activity.

### 3.2 Ovarian cancer microtissues develop cell line-specific histological architectures

Histological examination was performed to further characterize the architectural features of ovarian cancer microtissues during prolonged 3D culture. H&E staining at days 14 and 21 revealed distinct patterns of cellular organization along the collagen scaffold, consistent with the morphological differences observed by bright-field imaging (Fig. 2). Across the models, tumour cells predominantly localized to the collagen surface, forming multicellular layers and aggregates with varying degrees of organization.

**Figure 2.**
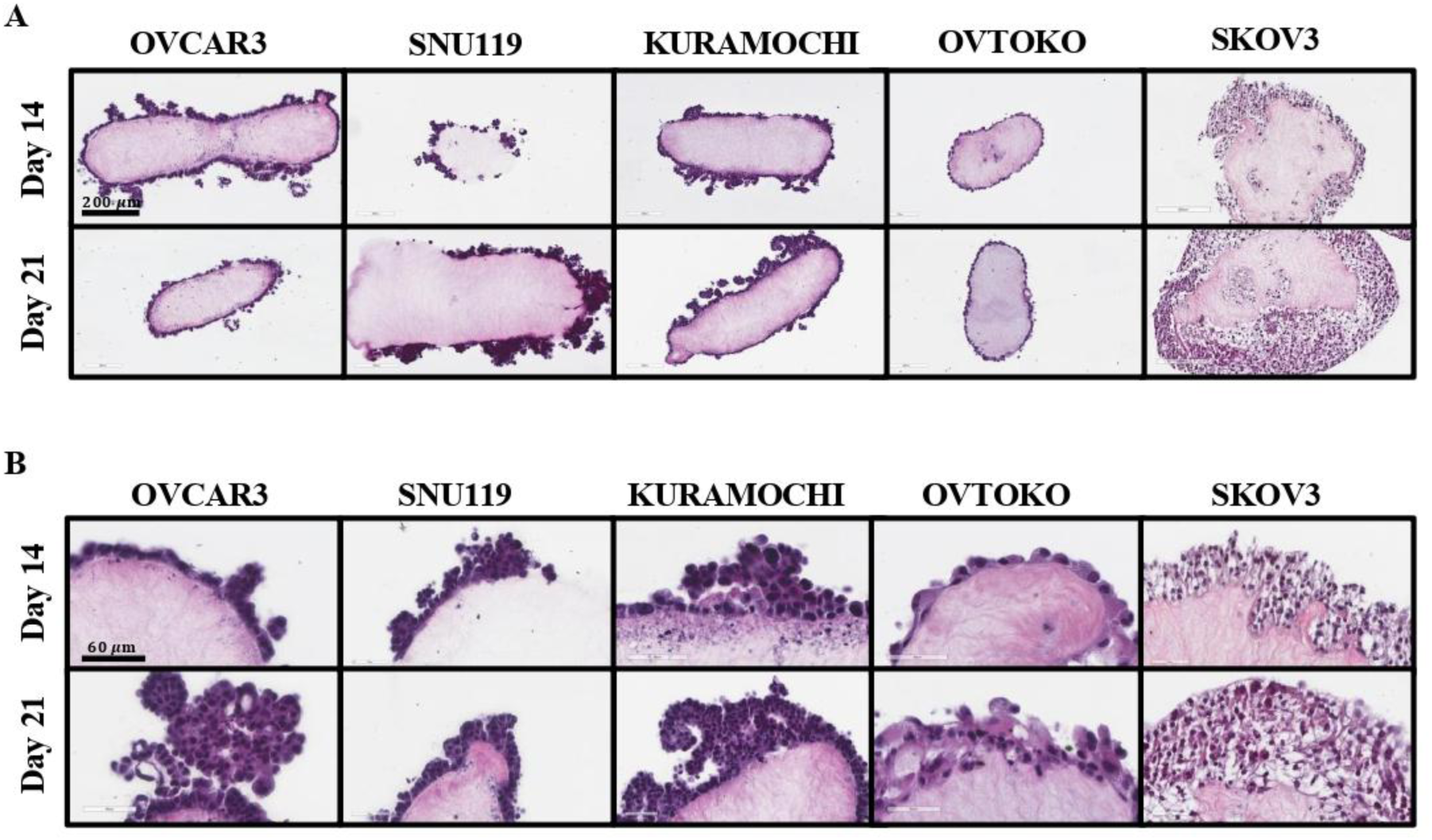
Histological characterization of ovarian cancer microtissues cultured in a 3D collagen-based matrix. Representative H&E-stained sections of ovarian cancer microtissues generated from five ovarian cancer cell lines (OVCAR3, SNU119, KURAMOCHI, OVTOKO, and SKOV3) at days 14 and 21 of culture. **(A)** Low-magnification (10X) images illustrating the overall histological architecture and distribution of tumour cells along the collagen scaffold. **(B)** Higher-magnification (40X) images highlighting cell line-dependent differences in cellular morphology and tissue organization, including papillary-like projections in OVCAR3 and SNU119, cohesive multicellular aggregates in KURAMOCHI, peripheral cellular organization with hobnail-like morphology in OVTOKO, and dense, irregular cellular arrangements in SKOV3. Scale bars = 200 µm (A) and 60 µm (B).

OVCAR3 and SNU119 microtissues exhibited irregular cellular projections extending outward from the collagen surface, with higher-magnification images revealing multicellular arrangements resembling papillary structures. These features were particularly evident at day 21, when more extensive cellular aggregation was observed. Kuramochi microtissues displayed a predominantly cohesive pattern of growth along the collagen boundary, with localized multicellular aggregates and less prominent papillary architecture. In contrast, OVTOKO exhibited a relatively thin peripheral layer of tumour cells, with individual cells displaying hobnail-like morphology. By day 21, OVTOKO cells had formed a thin, continuous multicellular layer covering the collagen surface. SKOV3 showed a distinct growth pattern characterized by extensive cellular accumulation along the collagen surface, forming a comparatively dense and irregular peripheral layer. Notably, despite the absence of an apparent increase in overall microtissue size, H&E sections obtained at day 14 and 21 revealed SKOV3 cells extending beneath the collagen surface and into the underlying scaffold, suggesting localized cellular invasion into the collagen matrix (Fig. 2).

These findings demonstrate that the collagen scaffold supports the development of distinct histological architectures across ovarian cancer models, with cell line-dependent differences in cellular distribution, aggregation, and surface organization that remained evident during prolonged 3D culture (Fig. 2).

### 3.3 Ovarian cancer microtissues retain cell line-associated immunophenotypic features

IHC analysis was performed to characterize the expression of ovarian cancer-associated markers in 3D microtissues at days 14 and 21. Representative staining patterns for PAX8, WT1, and p53 are presented in Fig. 3, while the corresponding quantitative and qualitative findings are summarized in Table 1. Nuclear PAX8 positivity was observed in 100% of tumour cells across all four evaluated models, including OVCAR3, SNU119, KURAMOCHI, and OVTOKO, at both time points, indicating the maintenance of a Müllerian-associated immunophenotype throughout prolonged culture. WT1 expression exhibited marked cell line-dependent heterogeneity but remained relatively stable between days 14 and 21. OVCAR3 and KURAMOCHI showed high WT1 positivity (approximately 70–90%), whereas SNU119 exhibited substantially lower expression (5–10%) and OVTOKO remained WT1-negative at both time points. Similarly, aberrant p53 staining patterns were maintained across all four models throughout the culture period. OVCAR3 exhibited an aberrant overexpression pattern, while KURAMOCHI, SNU119, and OVTOKO displayed staining patterns reported as aberrant mutant-type expression (Fig. 3; Table 1).

**Figure 3.**
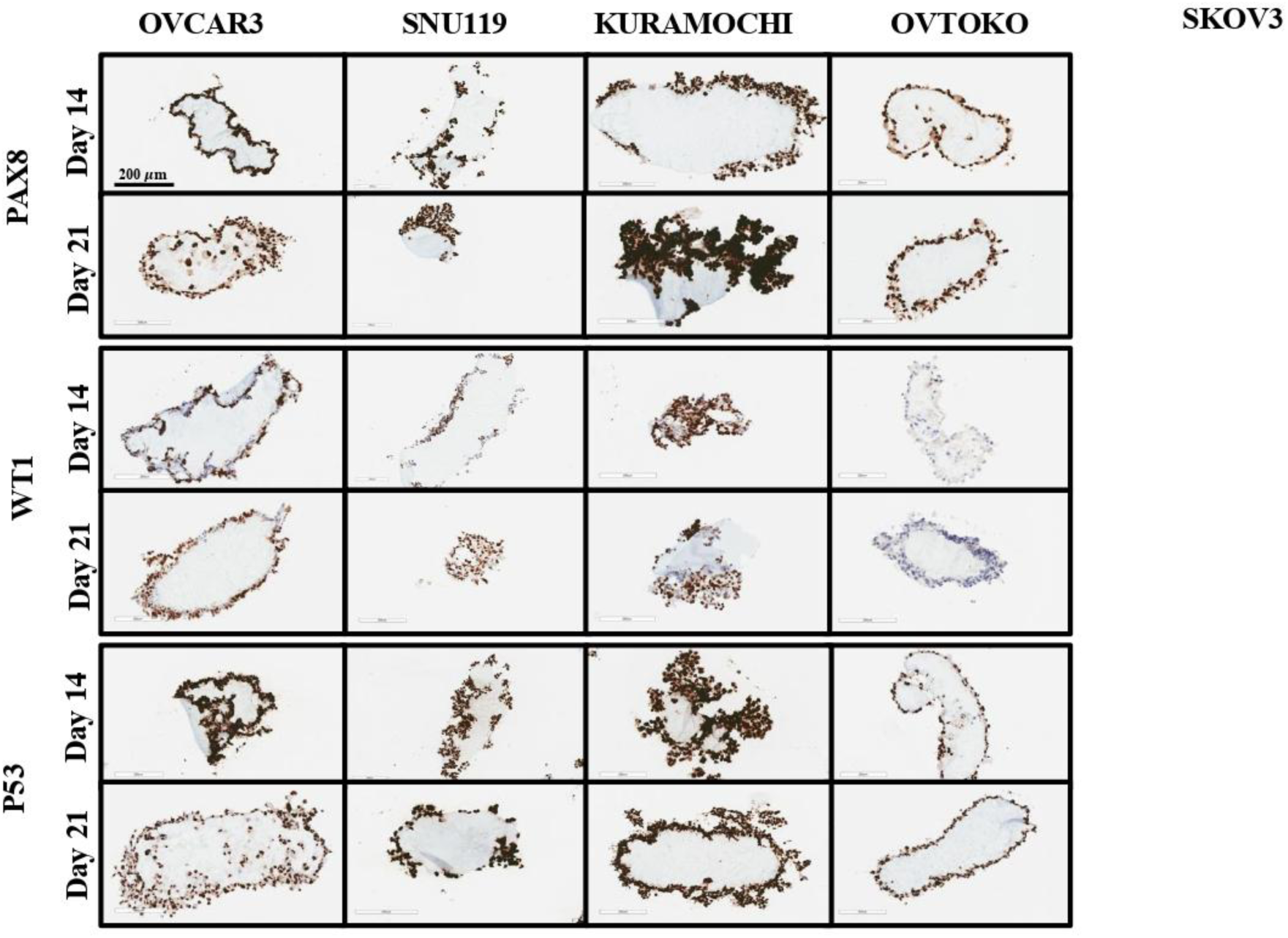
IHC characterization of ovarian cancer microtissues cultured in a 3D collagen matrix. Representative IHC staining of ovarian cancer microtissues generated from OVCAR3, SNU119, KURAMOCHI, and OVTOKO cell lines at days 14 and 21 of culture. **(A)** PAX8 staining demonstrating nuclear positivity across all four evaluated cell lines, consistent with the maintenance of a Müllerian-associated immunophenotype. **(B)** WT1 staining illustrating cell line-dependent differences in nuclear expression, with prominent positivity in OVCAR3 and KURAMOCHI, limited positivity in SNU119, and negative staining in OVTOKO. **(C)** p53 staining showing distinct nuclear staining patterns across the evaluated ovarian cancer models. Representative images illustrate the distribution and localization of these markers within the collagen-based microtissues. Scale bars = 200 µm,

**Table 1.** IHC characterization of ovarian cancer microtissues following 3D culture. Summary of IHC staining results for PAX8, WT1, and p53 in ovarian cancer microtissues generated from OVCAR3, SNU119, KURAMOCHI, and OVTOKO cell lines at days 14 and 21 of culture. Marker expression is reported according to the observed staining patterns and pathological assessments, highlighting cell line-dependent differences in ovarian cancer-associated immunophenotypic characteristics.

| Cell line | Time point | PAX8<br>(% positive) | WT1<br>(% positive) | p53 (expression pattern) |
| --- | --- | --- | --- | --- |
| OVCAR3 | Day 14 | 100 | ~80 | Aberrant (overexpression) |
|  | Day 21 | 100 | ~70 | Aberrant (overexpression) |
| KURAMOCHI | Day 14 | 100 | ~80 | Aberrant (mutant pattern) |
|  | Day 21 | 100 | ~90 | Aberrant (mutant pattern) |
| SNU119 | Day 14 | 100 | 5–10 | Aberrant (mutant pattern) |
|  | Day 21 | 100 | 5–10 | Aberrant (mutant pattern) |
| OVTOKO | Day 14 | 100 | 0 | Aberrant (mutant pattern) |
|  | Day 21 | 100 | 0 | Aberrant (mutant pattern) |

Collectively, these findings demonstrate that collagen-based 3D microtissues maintain distinct ovarian cancer-associated immunophenotypic profiles over 21 days, with minimal variation in the evaluated marker expression patterns between days 14 and 21.

### 3.4 Three-dimensional collagen culture alters therapeutic response compared with monolayer culture

To evaluate the influence of 3D culture on therapeutic response, ovarian cancer cells grown as collagen-based microtissues or conventional 2D monolayers were treated with increasing concentrations of carboplatin, paclitaxel, niraparib, and olaparib. Dose–response analysis revealed heterogeneous treatment responses across the five ovarian cancer models, with differences in drug sensitivity observed between 2D and 3D culture conditions (Fig. 4).

**Figure 4.**
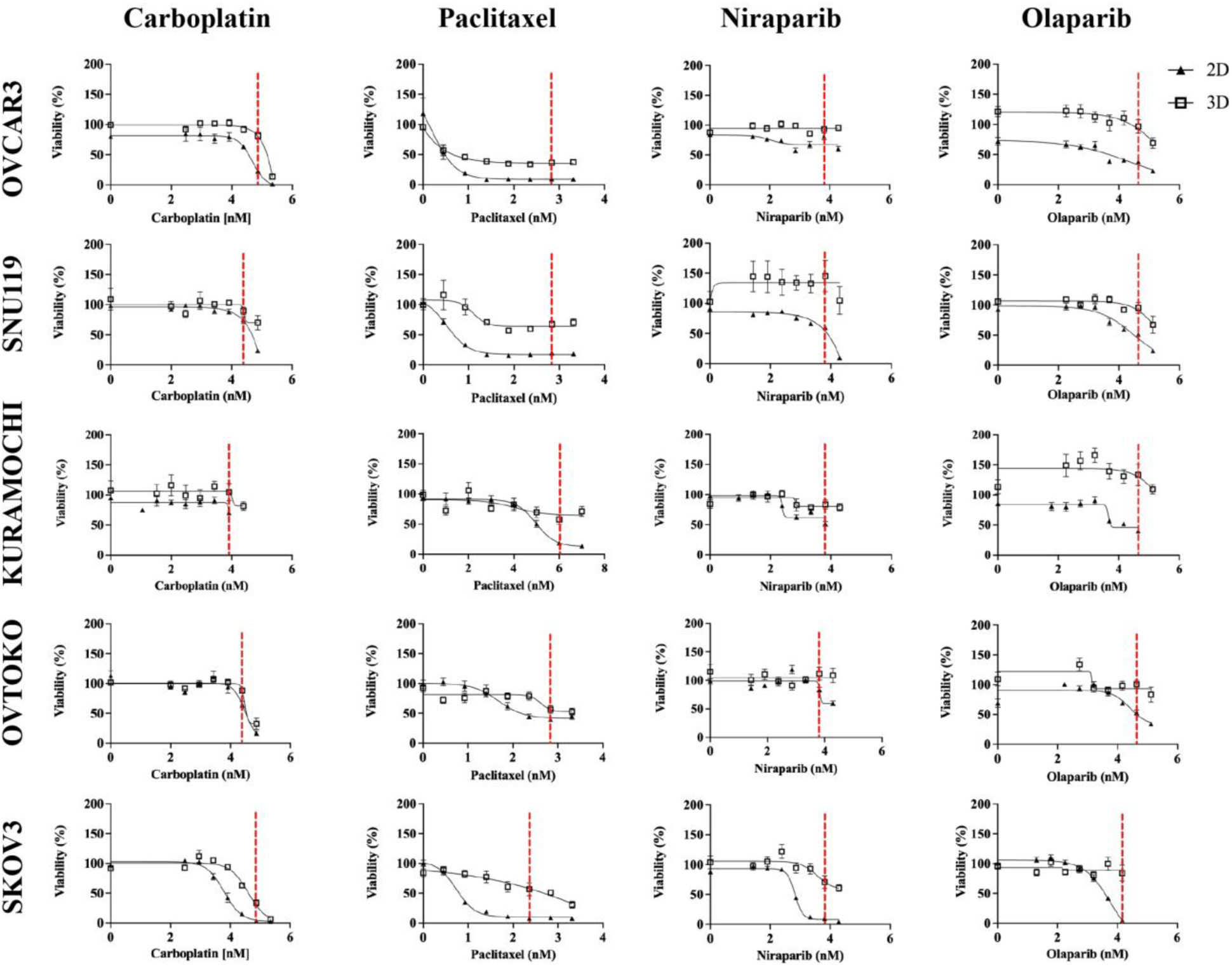
Comparative drug responses of ovarian cancer cell lines cultured under 2D and 3D conditions. Dose–response curves illustrating the effects of carboplatin, paclitaxel, niraparib, and olaparib on the viability of five ovarian cancer cell lines (OVCAR3, SNU119, KURAMOCHI, OVTOKO, and SKOV3) cultured as conventional 2D monolayers or collagen-based 3D microtissues. Cell viability (%) is plotted against drug concentration for each treatment, with 2D cultures represented by triangles and 3D cultures by squares. The curves illustrate cell line- and drug-dependent differences in therapeutic responses between the two culture conditions. Vertical red dashed lines indicate the maximum plasma concentration (Cmax) for each drug. Data are presented as mean ± standard error of the mean (SEM) from three independent experiments (n = 3). Dose–response curves were fitted using nonlinear regression with a variable-slope four-parameter logistic (4PL) model [log(inhibitor) versus response].

Responses to conventional chemotherapeutic agents varied across the models. Carboplatin treatment resulted in higher estimated IC50 values in 3D OVCAR3 and SKOV3 cultures, whereas SNU119 and KURAMOCHI exhibited lower estimated IC50 values relative to their 2D counterparts. OVTOKO showed comparatively similar carboplatin sensitivity under both culture conditions. Paclitaxel treatment also revealed divergent responses, with higher estimated IC50 values in 3D SNU119, OVTOKO, and SKOV3 cultures, while OVCAR3 and KURAMOCHI exhibited lower estimated values in 3D (Figs. 4, 5).

PARP inhibitor treatment further highlighted differences in therapeutic response among the models. Niraparib was associated with higher estimated IC50 values in 3D KURAMOCHI and SKOV3 cultures, whereas OVCAR3 and SNU119 exhibited lower estimated values. Similarly, olaparib treatment resulted in higher estimated IC50 values in OVCAR3 and KURAMOCHI microtissues, while SNU119 and OVTOKO showed lower estimated IC50 values compared with their corresponding 2D cultures. These differences were further illustrated by the log10-transformed 3D/2D IC50 ratios, which varied considerably across the evaluated drug–cell line combinations (Fig. 5).

**Figure 5.**
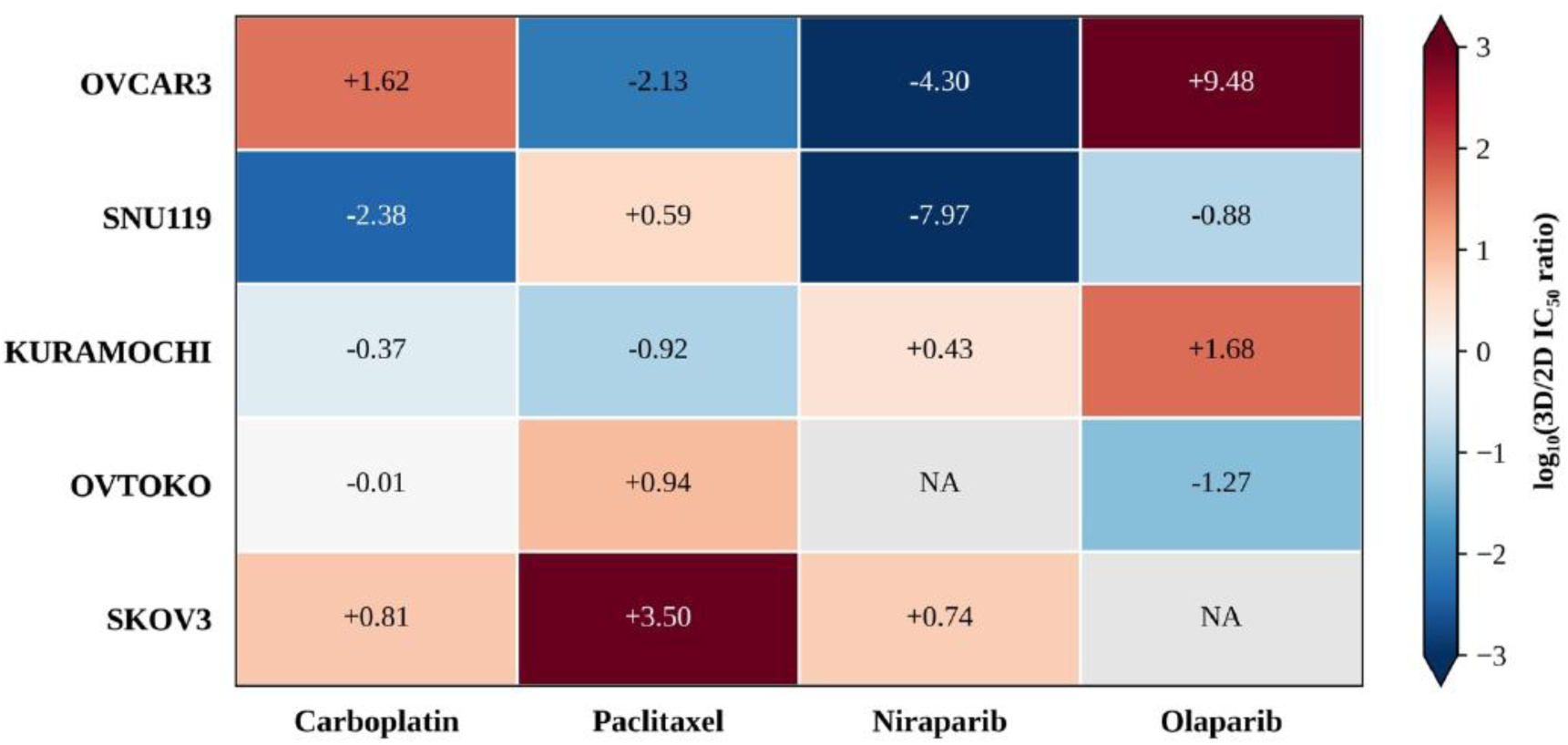
Heatmap visualization of relative drug sensitivity in 3D versus 2D ovarian cancer culture models. Heatmap illustrating the log₁₀-transformed ratios of half-maximal inhibitory concentrations (IC₅₀) between 3D and 2D cultures for five ovarian cancer cell lines (OVCAR3, SNU119, KURAMOCHI, OVTOKO, and SKOV3) treated with carboplatin, paclitaxel, niraparib, and olaparib. Values were calculated as log₁₀ (IC₅₀,3D/IC₅₀,2D). Positive values (red) indicate higher estimated IC₅₀ values in 3D cultures, corresponding to reduced apparent drug sensitivity relative to 2D cultures, whereas negative values (blue) indicate lower estimated IC₅₀ values in 3D cultures, corresponding to increased apparent drug sensitivity. Values approaching zero (white) indicate comparable estimated IC₅₀ values between the two culture conditions. The colour scale was limited to −3 and +3 for visualization, while numerical annotations display the uncapped log₁₀-transformed ratios. Grey cells labelled NA indicate unavailable or non-stable IC₅₀ ratios.

Collectively, these findings indicate that collagen-based 3D culture is associated with heterogeneous therapeutic responses rather than a uniform increase in drug resistance. The direction and magnitude of the estimated IC50 shifts varied according to both cell line and therapeutic agent, highlighting the distinct functional responses of ovarian cancer models under 3D culture conditions.

## 4. Discussion

In this study, we established a collagen-based 3D ovarian cancer microtissue model using five epithelial ovarian cancer cell lines representing distinct histological backgrounds, including the serous-associated models OVCAR3, KURAMOCHI, and SNU119, the ovarian clear cell carcinoma model OVTOKO, and non-HGSOC adenocarcinoma model SKOV3 ^(30)^. The models was comprehensively evaluated through bright-field and immunofluorescence imaging, histological examination, IHC characterization, and comparison of therapeutic responses between 3D microtissues and conventional 2D cultures. Collectively, our findings demonstrate that the collagen scaffold supported long-term tumour cell viability, maintained distinct histological architectures and immunophenotypic characteristics, and generated drug-response profiles that differed from corresponding monolayer cultures, highlighting its potential as a physiologically relevant in vitro model of ovarian cancer.

The successful establishment of long-term microtissues across all five ovarian cancer models demonstrates the suitability of type I collagen as a scaffold for 3D ovarian cancer culture. Type I collagen is one of the principal structural components of the ovarian tumour extracellular matrix and contributes to tumour progression through both biochemical and mechanical signalling rather than serving solely as structural support ^(15, 31)^. Unlike rigid plastic culture surfaces, collagen provides a matrix environment that supports cell–matrix interactions involved in adhesion, proliferation, migration, differentiation, mechanotransduction, and therapeutic response ^(14)^. Consistent with these biological functions, all cell lines remained viable throughout the 21-day culture period. Longitudinal CellTiter-Glo 3D measurements revealed distinct cell line-dependent changes in ATP-associated metabolic activity, with increasing luminescence signals in OVCAR3 and SNU119, declining signals in KURAMOCHI and OVTOKO, and a transient increase followed by a reduction in SKOV3 (Fig. 1C). However, both confocal live/dead imaging at days 14 and 21 (Fig. 1B) and H&E staining (Fig. 2) provided complementary spatial evidence of viable cells associated with the collagen microtissues with minimal cell death. Ovarian cancer cells exhibit metabolic plasticity, allowing them to adapt their metabolism energy by altering their reliance on glycolysis, mitochondrial oxidative phosphorylation, and fatty acid oxidation in response to changes in the tumour microenvironment ^(32)^. Previous studies have demonstrated that ovarian cancer cells can utilize aerobic glycolysis, characterized by lactate production, as well as fatty acid metabolism to meet their energy demands ^(33)^. These metabolic adaptations may contribute to the differences in ATP-associated metabolic activity observed among our ovarian cancer microtissue models ^(34)^. However, further studies are required to characterize the underlying metabolic pathways and determine their contributions to the cell line-dependent differences observed during 3D culture. Regardless, these observations indicate that the collagen scaffold maintained long-term cell viability while supporting distinct patterns of tumour cell organization during prolonged 3D culture (Fig. 1A–C).

Beyond supporting long-term survival, the collagen scaffold permitted the emergence of distinct architectural growth patterns among the ovarian cancer models. Bright-field imaging at days 14 and 21 showed that OVCAR3 and SNU119 developed prominent papillary-like projections extending from the microtissue surface, whereas KURAMOCHI displayed a predominantly clumpy surface-spreading growth pattern with heterogeneous cellular organization along the collagen scaffold. In contrast, OVTOKO exhibited a relatively compact morphology with a smoother and more uniform surface contour, while SKOV3 demonstrated extensive cellular accumulation and a comparatively compact but irregular microtissue morphology (Fig. 1A). These distinct morphologies became increasingly apparent during prolonged culture, demonstrating that the collagen microenvironment did not impose a uniform pattern of growth across the different models. Rather, each cell line established a distinct 3D architecture, suggesting that the collagen scaffold provides a permissive environment in which intrinsic, cell line-specific growth patterns can emerge. Histological examination further supported the architectural differences observed by bright-field microscopy and revealed that selected features of the microtissues resembled those found in intact ovarian tumours ^(35)^. Most notably, H&E sections confirmed that the projections observed in OVCAR3 and SNU119 consisted of multicellular structures extending outward from the collagen surface and exhibiting a papillary-like organization (Fig. 2). This finding is particularly relevant to their serous-associated phenotype, as papillary and micropapillary growth represent well-recognized architectural patterns of serous ovarian carcinomas ^(36)^. In contrast, KURAMOCHI displayed predominantly cohesive multicellular aggregates along the collagen boundary with substantially less prominent papillary organization, illustrating architectural heterogeneity even among serous-associated models. OVTOKO exhibited a distinct histological pattern characterized by a relatively thin peripheral distribution of tumour cells along the collagen surface with hobnail-like cellular morphology, a feature associated with ovarian clear-cell carcinoma ^(37)^ (Fig. 2). Importantly, the predominantly surface-associated organization observed across the models is relevant to ovarian cancer dissemination, during which tumour cells attach to and establish deposits along peritoneal and organ surfaces. Although the present system does not reproduce the complete peritoneal microenvironment as it lacks several important cell types such as mesothelial cells, it provides an accessible collagen surface that supports this aspect of tumour organization.

The histotype-associated architectural features observed by bright-field imaging and H&E were further supported by IHC profiling, which demonstrated the maintenance of selected ovarian cancer-associated immunophenotypic characteristics. All evaluated microtissue models exhibited nuclear PAX8 positivity at both days 14 and 21, consistent with a Müllerian-associated immunophenotype ^(38)^ (Fig. 3; Table 1). Despite culture in the same collagen formulation, the models retained distinct immunophenotypic profiles. OVCAR3 and KURAMOCHI maintained relatively high WT1 expression, whereas SNU119 exhibited substantially lower positivity and OVTOKO remained WT1-negative at both time points. Aberrant p53 staining patterns were also observed across the four models, with an overexpression pattern specifically reported for OVCAR3 (Fig. 3; Table 1). These findings are broadly consistent with the distinct immunophenotypic characteristics associated with serous and clear-cell ovarian carcinomas ^(39–41)^. When considered together with the papillary-like architecture observed in OVCAR3 and SNU119 and the distinct growth patterns of the other models, these findings demonstrate that the collagen microtissue system maintains selected histotype-associated features spanning tissue architecture and immunophenotype. Thus, rather than reproducing only the 3D shape of a tumour, the model captures selected structural and immunophenotypic characteristics associated with ovarian cancer, while maintaining the inter-model heterogeneity that is a defining feature of the disease.

Having established that the collagen microtissues maintain distinct structural features and ovarian cancer-associated IHC marker expression patterns, we next examined whether these characteristics were accompanied by altered therapeutic responses. Across the four agents tested, the influence of 3D culture on drug sensitivity varied considerably among cell lines and treatments, rather than producing a uniform increase in therapeutic resistance (Figs. 4 and 5). Importantly, this heterogeneity indicates that the effects associated with collagen-based culture depend on both the cellular model and the therapeutic agent. For carboplatin, OVCAR3 and SKOV3 exhibited higher estimated IC50 values in 3D than in 2D, whereas SNU119 and KURAMOCHI showed lower estimated values, and OVTOKO displayed a comparatively similar response between culture conditions (Figs. 4 and 5). Previous studies have demonstrated that extracellular matrix composition and stiffness can influence platinum resistance in ovarian cancer through integrin-mediated adhesion, focal adhesion kinase (FAK), and YAP-associated mechanotransduction pathways ^(42)^. These findings suggest that collagen-associated signalling may contribute to the differential carboplatin responses observed in our models, although these pathways were not directly assessed. Paclitaxel also produced divergent responses, with higher estimated IC50 values in 3D SNU119, OVTOKO, and SKOV3 cultures and lower estimated values in OVCAR3 and KURAMOCHI (Figs. 4 and 5). The pronounced reduction in apparent paclitaxel sensitivity in 3D SKOV3 is particularly interesting given previous evidence linking collagen I culture to altered microtubule-associated signalling. Specifically, 3D collagen I culture has been reported to increase expression of the microtubule-associated protein tau in ovarian cancer cells, while tau silencing enhanced paclitaxel sensitivity in collagen-cultured SKOV3 cells ^(43)^. Furthermore, altered integrin-dependent adhesion and FAK signalling have been associated with increased microtubule dynamics and paclitaxel resistance in ovarian cancer models ^(44)^. These findings raise the possibility that collagen-associated changes in cell adhesion and microtubule regulation may contribute to the differential paclitaxel responses observed here. However, these mechanisms were not directly investigated in the present study, and their potential contributions remain to be established. PARP inhibition similarly revealed model-dependent responses. Niraparib treatment was associated with higher estimated IC50 values in 3D Kuramochi and SKOV3 cultures but lower values in OVCAR3 and SNU119. Olaparib treatment produced higher estimated IC50 values in 3D OVCAR3 and KURAMOCHI cultures, whereas SNU119 and OVTOKO exhibited lower estimated values relative to their 2D counterparts (Figs. 4 and 5). These findings are particularly noteworthy when considered alongside the previously reported BRCA1/2 characteristics of ovarian cancer cell lines. KURAMOCHI carries a reported deleterious BRCA2 nonsense mutation (c.6952C>T; p.Arg2318Ter), which would be expected to increase susceptibility to PARP inhibition through impaired homologous recombination repair ^(45)^. Nevertheless, our results showed higher estimated olaparib and niraparib IC50 values in 3D KURAMOCHI microtissues than in their 2D counterparts. This observation is consistent with previous evidence that collagen I can contribute to PARP inhibitor resistance in ovarian cancer models, suggesting that extracellular matrix-associated survival or DNA damage response signalling may modify therapeutic sensitivity even in BRCA2-mutant cells ^(45)^. OVCAR3 provides another interesting example. Although this cell line has not consistently been characterized as carrying a deleterious BRCA1/2 mutation, previous functional studies identified impaired homologous recombination repair and sensitivity to the PARP inhibitor rucaparib ^(46)^. In the present study, OVCAR3 exhibited a higher estimated olaparib IC50 in 3D culture, suggesting that PARP inhibitor response may be influenced by factors beyond BRCA mutation status alone. These findings emphasize the complexity of therapeutic responses in ovarian cancer and highlight the potential influence of the 3D microenvironment on drug sensitivity. However, PARP inhibitor sensitivity alone cannot establish BRCA mutation status or homologous recombination deficiency, which would require independent molecular characterization (40).

Collectively, the drug-response experiments provide functional support for the preceding morphological, histological, and immunophenotypic characterization of the model. The same microtissues that developed distinct architectures and retained selected histotype-associated features also exhibited heterogeneous therapeutic responses, with the direction and magnitude of estimated IC50 changes varying across cell lines and treatments. Thus, the collagen platform captures complementary structural, immunophenotypic, and functional dimensions of ovarian tumour biology. Although it does not reproduce the full complexity of the tumour microenvironment, including stromal and immune components, vascular delivery, and systemic influences, its ability to combine long-term viability, surface-associated tumour architecture, immunophenotypic characteristics, and heterogeneous therapeutic responses within a single platform supports its utility as a complementary experimental model for investigating ovarian cancer biology and therapeutic response ^(47)^.

## 5. Conclusion

In this study, we developed a type I collagen-based 3D microtissue platform capable of supporting long-term viability across diverse ovarian cancer models. Despite being maintained within the same collagen environment, the models developed distinct patterns of surface-associated tumour organization and retained differences in tissue architecture and immunophenotype. These differences were also reflected functionally, as the microtissues displayed heterogeneous responses to chemotherapy and PARP inhibition. Importantly, the influence of 3D culture on drug sensitivity varied across cell lines and treatments rather than producing a uniform increase in resistance, highlighting the context-dependent relationship between collagen-based culture and therapeutic response. Together, these findings demonstrate that the collagen microtissue platform captures structural, immunophenotypic, and functional heterogeneity within a controlled 3D environment. Although the system does not reproduce the full complexity of the native tumour microenvironment, it supports several features relevant to ovarian cancer biology that are not readily represented in conventional monolayer culture. The platform therefore provides a tractable model for investigating ovarian cancer biology, tumour– matrix interactions, and therapeutic response, with potential for future incorporation of additional stromal and immune components to further increase its physiological relevance.

